# Single-cell data and correlation analysis support the independent double adder model in both *Escherichia coli* and *Bacillus subtilis*

**DOI:** 10.1101/2020.10.06.315820

**Authors:** Guillaume Le Treut, Fangwei Si, Dongyang Li, Suckjoon Jun

## Abstract

The reference point for cell-size control in the cell cycle is a fundamental biological question. We previously reported that we were unable to reproduce the conclusions of Witz *et al.*’s *eLife* paper (Witz, van Nimwegen, and Julou 2019) entitled, “Initiation of chromosome replication controls both division and replication cycles in *E. coli* through a double-adder mechanism”, despite extensive efforts. In this ‘replication double adder’ (RDA) model, both replication and division cycles are determined via replication initiation as the sole implementation point of size control. Witz *et al.* justified the RDA model using a type of correlation analysis (the “***I***-value analysis”) that they developed. By contrast, we previously showed that, in both *Escherichia coli* and *Bacillus subtilis,* replication initiation and cell division are determined by balanced biosynthesis of key cell cycle proteins (e.g., DnaA for initiation and FtsZ for cell division) and their accumulation to their respective threshold numbers, which Witz *et al.* coined the ‘independent double adder’ (IDA) model. The adder phenotype is a natural quantitative consequence of these mechanistic principles. In a recent *bioRxiv* response to our report, Witz and colleagues explicitly confirmed two important limitations of the ***I***-value analysis: (1) it is only applicable to non-overlapping cell cycles, wherein *E. coli* is known to deviate from the adder principle, and (2) it is only applicable to select biological models and, for example, cannot evaluate the IDA model. These limitations of the ***I***-value analysis were not explained in the original *eLife* paper and were overlooked during the review process. In this report, we show using data analysis, mathematical modeling, and experiments why the ***I***-value analysis - in its current implementation - cannot compare different biological models. Furthermore, the RDA model is incompatible with the adder principle and is not broadly supported by experimental data. For completeness, we also provide a detailed point-by-point response to Witz *et al.*’s response (Witz, Julou, and van Nimwegen 2020) in the Supplemental Information.

## Introduction

Papers in interdisciplinary research can be difficult to evaluate as they connect multiple disciplines. As such, the reader sometimes either must spend a significant effort to understand the details of the paper, or should take a leap of faith. Last year, we found ourselves spending an unusually large number of hours trying to understand a paper from Witz *et al*. that first appeared in *bioRxiv* (Witz, van Nimwegen, and Julou 2019), and communicated our questions and concerns with the authors. That paper was eventually published in *eLife* with minor changes. As their replication double adder (RDA) model of cell-size control contradicted the biological evidence accumulated in our lab over a decade, we decided to reproduce their results by directly applying their own ***I***-value analysis to both Witz *et al*.’s data as well as our data. We also conducted new experiments as needed over the course of several months. Furthermore, we derived analytical results on the RDA model.

We made our results publicly available in the form of a comment piece on Witz *et al. eLife* article (Le Treut et al. 2020). Witz and colleagues responded to our comment via *bioRxiv* (Witz, Julou, and van Nimwegen 2020). The most surprising additional information in their response was that (1) the ***I***-value analysis, in fact, is only applicable to *E. coli* with non-overlapping cell cycles, (2) by construction, the ***I***-value analysis is only applicable to select biological models, and (3) one of those models the ***I***-value analysis cannot evaluate is the ‘independent double adder’ (IDA) model, even for non-overlapping cell cycles.

These limitations are unfortunate since the hallmark of *E. coli* cell cycle is its multifork replication. For example, the Helmstetter-Cooper model was directly motivated by the existence of overlapping cell cycles. Furthermore, the celebrated (nutrient) growth law (Schaechter, Maaløe, and Kjeldgaard 1958) is also applicable to the “fast” growth conditions wherein cell cycles typically overlap. By contrast, *E. coli* does not strictly follow the adder principle in slow-growing cells as reported by Wallden et al. (2016) and explained by Si et al. (2019). Therefore, the clarification by Witz, Julou, and van Nimwegen (2020) raises other questions such as whether the RDA model is also intended for overlapping cell cycles, or whether it is still limited to non-overlapping cell cycles as the ***I***-value analysis itself. These limitations were not explained in their original *eLife* paper, and were overlooked by the referees during the review process (eLife 2019).

After careful examination of their response and re-examination of our own analysis, we stand by the conclusions reached in our previous comment (Le Treut et al. 2020). Specifically, we show below that (1) The ***I***-value analysis cannot compare different biological models or predict biologically correct models, and (2) The RDA model is not broadly supported by extensive data from two evolutionary distant model organisms and cannot be a sound model for *E. coli* cell-cycle and cell-size control.

This focused comment piece for *bioRxiv* is to address all major points in Witz *et al.*’s original paper (Witz, van Nimwegen, and Julou 2019) and their response (Witz, Julou, and van Nimwegen 2020), and help the community understand key subtle issues in drawing biological conclusions from statistical analysis of high-throughput single-cell data. The readers are thus encouraged to read the original *eLife* paper (Witz, van Nimwegen, and Julou 2019), our previous comment (Le Treut et al. 2020), their response (Witz, Julou, and van Nimwegen 2020), and this report. Since we view our exchanges as an attempt to resolve scientific disagreements, we set an internal rule to address only the scientific issues.

## Results

### 1. The ‘replication double adder’ vs. the ‘independent adder’ models, and what motivated our analysis

In their response via bioRxiv, Witz *et al.* defined, coined, and compared two models, the ‘replication double adder’ (RDA) vs. the ‘independent adder’ (IDA). For consistency, we will follow their conventions in this response. These two models are illustrated in Figure 1a, and each model can be characterized by a set of two correlations (Figure 1b).

**Figure 1.**
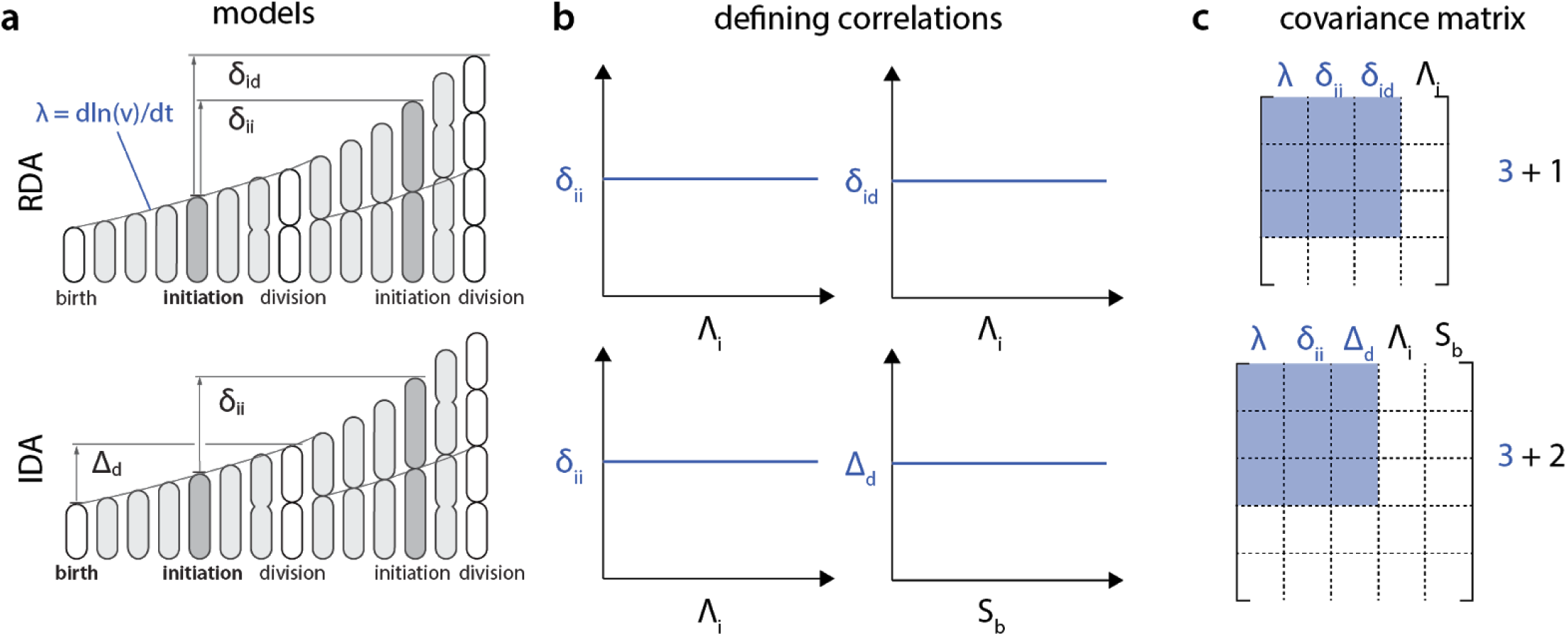
**a.** Two models defined in (Witz, Julou, and van Nimwegen 2020): Replication double adder (RDA; top) and Independent double adder (IDA; bottom). Each model can be described by one “temporal” parameter, the growth rate λ = 1/v dv/dt (where v is the volume of the cell during growth), and two “spatial” parameters. For the RDA model, the reference point is the initiation mass Λ_i_, with added size from initiation to determine division (δ_id_) and next initiation (δ_ii_). For the IDA model, these parameters are λ, δ_ii_, and Λ_d_ as illustrated. **b**. Each of these models is characterized by a set of two correlations, or the lack thereof. Note that these defining correlations require 1 additional parameter Λ_i_ for the RDA model, whereas the IDA model requires 2 additional parameters, Λ_i_ and S_b_. **c**. As a result, the covariance matrix for the ***I***-value analysis of these models, according to Witz *et al.*, would be 4×4 for the RDA model and 5×5 for the IDA model. Therefore, the ***I***-values of these models cannot be meaningfully compared.

The most important biological differences between the two models are about which stage of the cell cycle cell-size control is imposed. The sole implementation point of the RDA model is the initiation of DNA replication. The model then suggests that the cell measures two separate added sizes: from initiation to initiation δ_ii_, and from initiation to division δ_id_, with the initiation mass per origin of replication Λ_i_ as the reference point. Therefore, a full description of the RDA model requires 3 independent control parameters {δ_ii_, δ_id_, Λ}, including the growth rate Λ as a temporal variable. By contrast, the IDA model has two separate reference points: replication initiation (at size Λ_i_) and birth division (at size S_b_), with respective added sizes: from initiation to initiation δ_ii_, and from division to division Δ_d_. Therefore, the three control parameters that define the IDA model are {δ_ii_, Δ_d_, Λ}.

While these two models may look similar, their underlying biological mechanisms are fundamentally different and even incompatible with each other. In the RDA model, the cell monitors the size added since initiation and, when it reaches a critical value δ_id_ that is set independent of the cell size at initiation, the cell triggers division. It is unclear what mechanisms would be activated at replication initiation, and how they would trigger cell division. The problem becomes even more difficult to comprehend with overlapping cell cycles (i.e. multifork replication). We do not know whether the RDA model is also intended for overlapping cell cycles and, if yes, the model would need an explanation as to how the hypothetical length-measuring device is activated first at initiation, and then track the added size continuously, and finally trigger cell division when the cell has added a fixed added size since initiation, over the course of multiple generations. We are currently unaware of any such molecules or devices in any organism, and it is these biological implications of the RDA model that motivated us to understand what led Witz *et al.* to their conclusion.

For the IDA model, we explained the mechanistic principles at the molecular level, i.e., cells produce division proteins (e.g., FtsZ) at the same rate as the growth rate, and trigger constriction when they have accumulated the proteins to a threshold number (Si et al. 2019). For initiation, cells produce initiator proteins (e.g., DnaA) at the same rate as the growth rate, and trigger initiation when they have accumulated the proteins to a threshold number per replication origin. The two processes progress independently unless they are forced to interact with each other. For example, if replication termination is delayed abnormally beyond the expected division timing (e.g., due to a significant delay in initiation), then division must wait [see, for example, Figure S3D in (Si et al. 2019)]. On the other hand, we showed that moderate perturbations to the initiation timing do not affect division and vice versa. We made falsifiable predictions of the IDA model and tested them experimentally (Si et al. 2019).

Based on the evidence accumulated so far, we stand by our previous conclusion in Le Treut et al. (2020) as elaborated below.

### 2. The *I-value analysis* in its current form cannot compare different models

In their original article in *eLife,* Witz *et al.* presented a method whereby potential biological models were ranked according to so-called ***I***-values. This analysis was presented as a general quantitative method to identify the model most compatible with experimental data, based on its full correlation structure. From this analysis, Witz *et al*. drew general conclusions, reflected in their title: “Initiation of chromosome replication controls both division and replication cycles in *E. coli* through a double-adder mechanism”.

In their response to our comment, Witz *et al*. clarified that the role of the ***I***-value analysis is not to discover biological mechanisms from all possible models, but to compare the correlation structure among a specific class of models. Although only 3 independent control parameters are sufficient to model the *E. coli* cell cycle (Figure 1a), Witz *et al.* suggested that the analysis of 4-variable combinations is needed to incorporate characteristic correlations of the RDA model.

The ***I***-value analysis could be a useful correlation analysis, if the following caveats can be overcome.

#### 2.1. The *I*-value analysis cannot compare different biological models

To see this, let us consider the two models in Figure 1. The first model is the RDA that can be defined by the 3 parameters {δ_ii_ δ_id_, Λ} (Figure 1a, top). Witz *et al*. argue that the analysis of 3×3 matrices can show degeneracy, because the defining two correlations, namely (Λ_i_, δ_ii_) and (Λ_i_, δ_id_), require an additional variable Λ_i_ (Figure 1b, top). For example, {δ_ii_, δ_id_, Λ} and {δ_ii_ Λ_i_, Λ} would both lead to maximum ***I***-values although one cannot infer the RDA model from either one alone. For this reason, Witz *et al*. suggested, and we agree, that the correlation structure of at least the following 4 variables {δ_ii_, δ_id_, Λ, Λ} is required to characterize the RDA model with an ***I***-value.

The problem with the above procedure is that the size of the covariance matrix becomes model dependent, and thus the ***I***-value analysis cannot compare different models. For example, the IDA model would require 5 variables for the ***I***-value analysis, because of the two defining correlations (Λ_i_, δ_ii_) and (S_b_, Δ_d_) (Figure 1b, bottom). Therefore, in addition to the three independent control parameters {δ_ii_, Δ_d_, Λ}, the ***I***-value analysis would require two additional parameters from the defining correlations (Figure 1b, bottom), namely, the initiation mass Λ_i_ and the birth size S_b_. The resulting covariance matrices would then be 5×5, not 4×4. However, the ***I***-values obtained from covariance matrices of different sizes cannot be meaningfully compared. Therefore, if we follow Witz *et al*. ‘s reasoning, the ***I***-value analysis is fundamentally limited to compare a specific class of models.

#### 2.2.

With this caveat in mind, Witz and colleagues nonetheless insisted on performing a 4-variable ***I***-value analysis for the *E. coli* cell cycle rather than a 3-variable analysis (Witz, Julou, and van Nimwegen 2020). In fact, we had already performed the 4-variable analysis in Le Treut et al. 2020 and the results, as shown in Figure 2 below, have been available via Github (https://github.com/junlabucsd/DoubleAdderArticle). The conclusion from the analysis is nevertheless clear. Both the 3-variable analysis and the 4-variable ***I***-value analysis are consistent with each other (Figure 2, top), and the IDA model scores higher than the RDA model.

**Figure 2.**
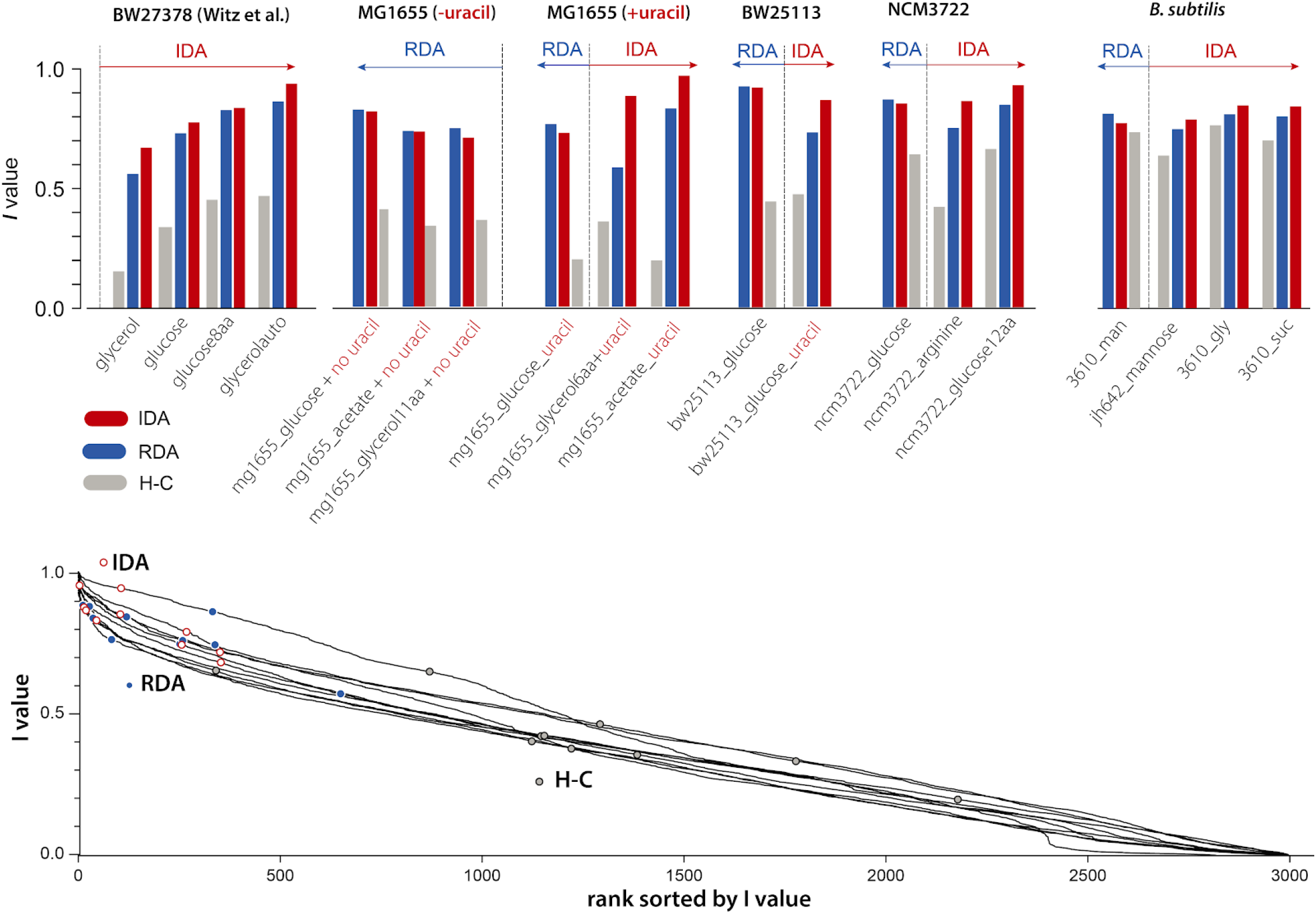
The 4-variable analysis of ***I***-values for the IDA, RDA and Helmstetter-Cooper models in different growth conditions in *E. coli* and *B. subtilis*. Most strikingly, the IDA model unanimously scores higher than the RDA model in Witz et al.’s data. In the MG1655 data, the RDA model initially scored marginally higher than the IDA model, but with the supplement of uracil the reversion of the order is apparent, as explained in our previous comment. The arrows in the figure show the set of data that favor the model indicated above the arrows. The bottom panel shows ***I***-values by decreasing order computed for all combinations of 4 variables among 18 cell cycle variables [see Figure S1 in (Le Treut et al. 2020)].

#### 2.3.

Many other combinations of variables can also produce even higher ***I***-values than both the RDA model and the IDA model (Figure 2, bottom). Therefore, it is not obvious how the ***I***-value analysis could be used to predict potential models of the cell cycle. Pruning biologically unsound models manually before applying the ***I***-value analysis would be daunting, since there are 3,060 possible combinations for the 4-variable analysis, 8,568 combinations for the 5-variable analysis, and so on.

## 3. The RDA model is analytically incompatible with the adder principle

In our comment, we presented a theoretical analysis showing that the RDA model is incompatible with the adder principle. In particular, we demonstrated that the RDA model predicts a negative slope when plotting 〈Δ_d_ | S_b_ 〉 versus S_b_, hence a deviation toward a sizer behavior. Specifically, we showed that in the RDA model, the mother/daughter correlation coefficient for division size reads:

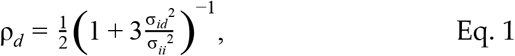

where σ_ii_^2^ and σ_id_^2^ are the variances of and δ_id_ respectively. The aforementioned slope is then 2 *ρ*_d_ −1 < 0.

Therefore, we were puzzled that simulations of the RDA model by Witz, van Nimwegen, and Julou 2019 reproduced the adder phenotype, which is not possible according to Eq. 1. We found that this agreement critically relied on the existence of fluctuations in the division ratio (or septum positioning). To show this, we ran simulations using the code included in Witz, van Nimwegen, and Julou (2019) by simply setting the variance in the division ratio to zero to simulate symmetric division. The results agreed with the prediction of Eq. 1 quantitatively.

In other words, the agreement of the simulations with the experimental adder in Witz, van Nimwegen, and Julou (2019) is not the properties of the RDA model, but the result of the fifth ad-hoc parameter, which was not included in the ***I***-value analysis of the RDA model. Furthermore, ad-hoc fluctuations can produce the adder-like phenotypes even when the underlying biological mechanisms are incompatible with the adder principle (Fig 2 in (Si et al. 2019); (Facchetti et al. 2019)). Therefore, to show the biological mechanisms, it is critical to make experimentally testable predictions beyond correlations based on the mechanistic hypothesis (Si et al. 2019).

As an additional note, following the standard practice in the open-source community, we made the code used for these simulations publicly available via GitHub (https://github.com/junlabucsd/DoubleAdderArticle). The code is version-controlled from Witz *et al.*’s own repository, and it should therefore be straightforward to check that we did not alter the code for RDA simulations in their repository.

In summary, the RDA model is analytically incompatible with the adder principle that is observed in most growth conditions for *E. coli* and *B. subtilis.*

## 4. The RDA model is based on one dataset, but the full datasets show different results

Witz and colleagues explained in their response to Le Treut et al. (2020) that they used the (Λ_i_, δ_id_) correlation to justify the RDA model and rule out the IDA model. Specifically, they stated that the IDA model predicts a “strong negative correlation”, whereas the RDA predicts a zero correlation. The following points are at odds with this claim.

1. The RDA model was derived from one experiment with non-overlapping cell cycle conditions. However, in these slow-growth conditions, *E. coli* is already known to deviate from the IDA phenotype, as shown before (Wallden et al. 2016). Si & Le Treut *et al.* specifically explained the observation mechanistically that the adder principle breaks due to active degradation of FtsZ in the slow growth conditions (Sekar et al. 2018; Männik, Walker, and Männik 2018; Si et al. 2019). They also demonstrated experimentally how to restore the adder phenotype by knocking out *clpX.*
2. The data itself shows negative correlations. For example, while the condition used in Witz, van Nimwegen, and Julou 2019 for the RDA model (Figure 3a) does not exhibit a clear trend, their three other experiments (Figure 3b) as well as most of our experimental data also show negative correlations (Figures 3c and 3d). However, even in the IDA model, the correlation should eventually vanish if the number of overlapping cell cycles increases significantly, as the initiation point and the cell division coupled with the initiation is separated by several generations. See the Supplemental Information for the analytical expression of the correlation for the IDA model.
3. Witz and colleagues emphasize the importance of comparing the full correlation structure of different models to data. Therefore, ruling out other biological models based on one correlation is self-contradictory.

**Figure 3.**
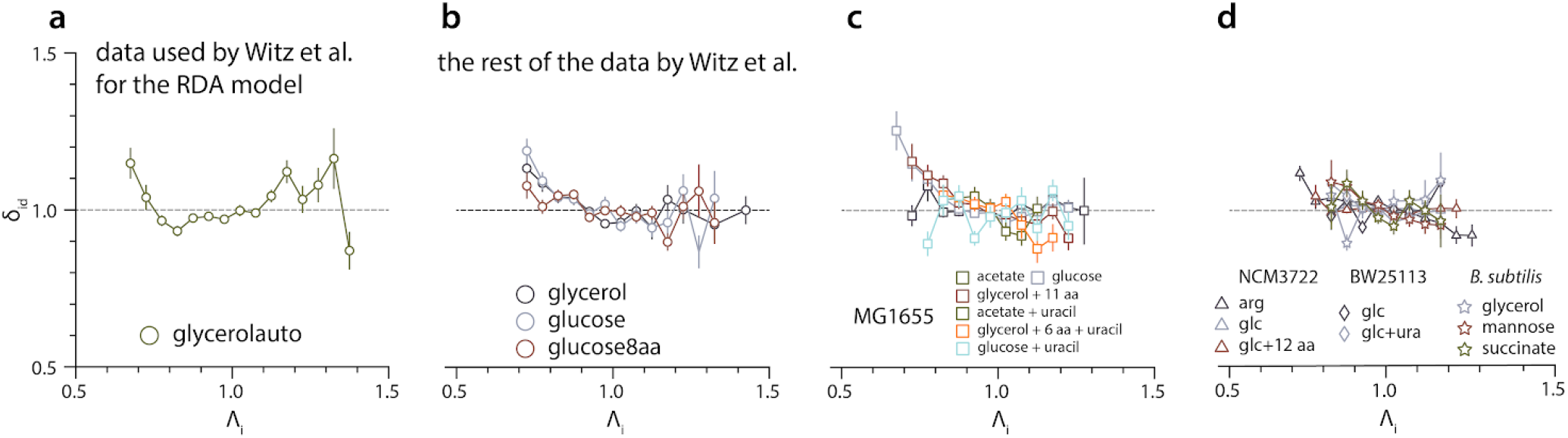
**a**. Witz et al.’s RDA model is exclusively based on this data set. **b-d**. 3 out of the 4 datasets of Witz et al. present a slight negative correlation. Most of the experimental data we obtained with *E. coli* MG1655, NCM3722, BW25113 and *B. subtilis* NCIB 3610 strain also show varying degrees of negative correlation, consistent with the IDA model.

## Materials and Methods

### Availability of analysis, results, and notes

All the numerical results discussed in the main text are available with detailed notes in the GitHub repository: https://github.com/junlabucsd/DoubleAdderArticle.

### Calculation of the (Λ_i_, δ_id_) correlation for the IDA model

In the IDA model, the cell size at division is determined by the cell size at birth S_b_ and the added size from birth to division Δ_d_; and the cell size per origin at initiation in the next cell cycle Λ_i_^(n+1)^ is determined by the added size per origin between consecutive replication initiation events δ_ii_, and the cell size per origin at initiation Λ_i_. The following relations hold:

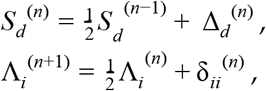

where the index *n* denotes the generation (or division cycle), and {Δ_d_^(n)^} and {Λ_i_^(n)^} are independently distributed random variables. Denoting *μ_dd_* = < Δ_*d*_ > and μ_*ii*_ = < δ_*ii*_ >, we have μ_*d*_ = < *S_d_* > = 2 μ_*dd*_ and μ_*i*_ = < Λ_*i*_, > = 2 μ_*ii*_. We also define the centered variables: 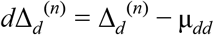, 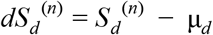, 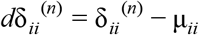 and 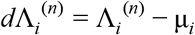. Denoting 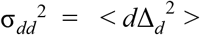 and 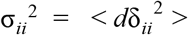, we have 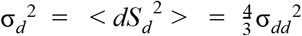 and 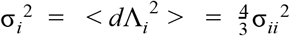.

The added size between initiation and division reads 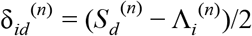, where the factor of 2 reflects the fact that replication origins double at initiation. Therefore:

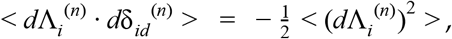

by independence of S_b_^(n)^ and Λ_i_^(n)^, and we obtain the Pearson correlation coefficient:

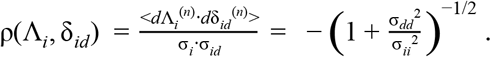

## Acknowledgments

We thank the members of the Jun lab and Hwa lab (UCSD) for helpful discussions, and many colleagues who generously shared their precious time for critical reading of this report. This work was supported by NIH grant R01 GM118565 (to S.J.) and NSF MCB-2016090.

## Supplemental Information

Below, we provide point-by-point responses to the comments raised by Witz, Julou, and van Nimwegen 2020.

### “Introduction”

*Le Treut et al. recently published a comment on our article Witz et al. (2019) and submitted the same comment for publication to eLife (Le Treut et al., 2020). The journal invited us to write a formal response and based on review of the editors and an external expert of both the comment of Le Treut et al. and our response, the journal decided to reject the comment of Le Treut et al. However, they strongly suggested we publish our response on bioRxiv to make sure it is available to the public.*

*The comment of Le Treut et al. severely criticized our work. In particular, Le Treut et al. claim that our data support a different model than the one we propose, that the correlation analysis that we introduced in any case cannot distinguish different models, and that our model is at odds with the ‘adder phenotype’ observed in the data. All these claims are false. As we will show below, the correlation analysis results that Le Treut et al. report are all meaningless because of basic errors in the application of this method. In addition, the claim that our model fails to reproduce the adder phenotype is based on comparing the experimental data not with our actual model, but with a mutilated version that the authors concocted by altering the simulation code that we provided.*

*Since all the results presented by Le Treut et al. are based on faulty analysis, we cannot see anything constructive in their contribution, and we fear that the main result of their comment will be to muddy the waters regarding our current understanding of the bacterial cell cycle. While data from the additional experiments that Le Treut et al. report might well provide a valuable addition to the literature, their analysis of this data is equally faulty, so that it is currently unclear what additional insights might be gained from this data.*

### “Division-centric? Replication-centric? What exactly are we talking about?

*The main impetus behind the comment of Le Treut et al. seems to have been a perceived contradiction between our results and the results they presented in Si et al. (2019), and much of their discussion concerns comparison of ‘their’ division-centric and ‘our’ replication-centric model of cell size homeostasis. However, in the absence of specifying concrete models, these terms are ambiguous and we suspect that many readers may easily lose track of what the actual differences are between these models, as well as to what extent the data presented in Si et al. (2019) and Witz et al. (2019) are even at odds or supporting one model over the other. We will thus start by giving a short overview of what precisely these models are and to what extent different data either support or reject them. Notably, given the large variety of cell cycle models that have been proposed in the literature, and the emphasis that Le Treut et al. put on contrasting their model with ours, the reader may be surprised to learn that these models are actually almost identical, differing in only one aspect. “*

The terms “division-centric” and “replication-centric” were introduced in (Witz, van Nimwegen, and Julou 2019), and we applied their terms and their models in our analysis.

*“To start from a historical reference point, the influential Helmstetter-Cooper model assumes that replication is initiated at a fixed critical cell size and that a fixed time elapses between replication initiation and division (Cooper and Helmstetter, 1968). Using a combination of microfluidics with time-lapse microscopy, in recent years researchers have been exploring to what extent the observed correlation structure of the fluctuations in the sizes and times at birth, initiation and division in single cells are consistent with such models of the cell cycle. In particular, if a model assumes that the cell cycle control mechanism acts through constraining a particular variable X, then one would expect single-cell fluctuations in X to be independent of fluctuations in other quantities. For example if, as the Helmstetter-Cooper model assumes, initiation is triggered when the cell reaches a critical size, one would expect fluctuations in the size at initiation to be independent of fluctuations in the size at birth. However, recent works, including ours (Fig. 2A), show that there is a clear positive correlation between size at birth and initiation (Micali et al., 2018a; Witz et al., 2019). **In this way, a particular model of the cell cycle can be falsified by showing that the observed correlations are at odds with predictions of the model.”***

Witz *et al.* emphasized in both their original paper and their response that their objective is to compare the full correlation structure, rather than ruling out using one correlation (see, for example, next page highlighted in yellow). Also importantly, the “positive correlation between size at birth and initiation” is only significantly seen in Witz *et al*.’s data (see Figure S1, left-most panel). By contrast, our data show zero to slightly positive correlations (see Figure S1, the other four panels).

**Figure S1.**
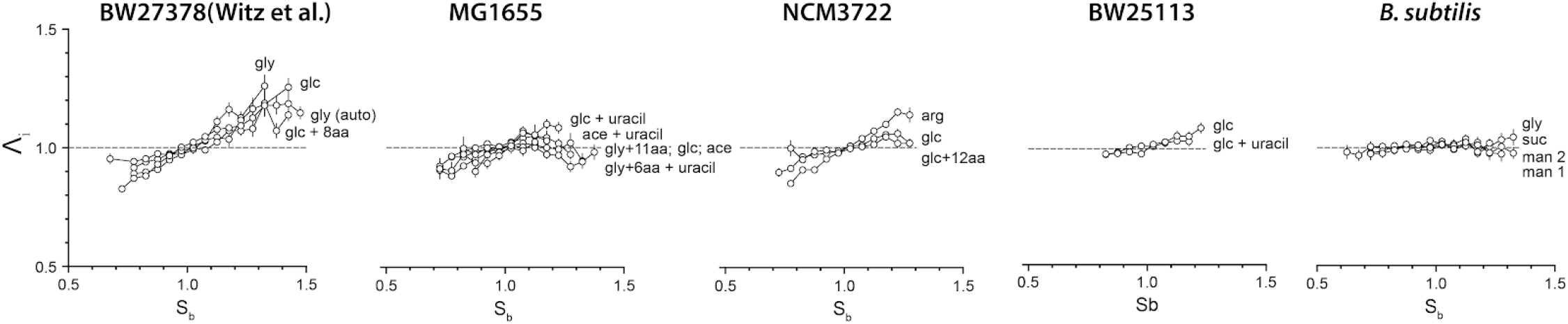
The (S_b_, Λ_i_) correlation between sizes at birth and initiation is only significant for Witz *et al*’s data. By contrast, most of the experimental data we obtained with *E. coli* MG1655, NCM3722, BW25113 and *B. subtilis* NCIB 3610 strain show zero to slightly positive correlations.

*“An alternative model of the replication cycle assumes replication initiation is controlled by an inter-initiation adder, i.e. that cells add a fixed volume per origin between consecutive replication events. Indeed, both in Si et al. (2019) and in our study (Witz, et al., 2019) it was observed that the added volume per origin fluctuates independently from the size at initiation, supporting this model.*

*However, it is important to note that this observed independence does not in itself prove that replication must be controlled by an inter-replication adder mechanism, i.e. other models may well also predict that there is no correlation between size at initiation and added volume between initiations.*

*As a case in point, the so called ‘adder phenotype’ corresponds to the observation that the volume added between birth and division is approximately uncorrelated with the cell’s size at birth (Amir, 2014; Osella et al., 2014; Campos et al., 2014; Taheri-Araghi et al., 2015). Although this observation can of course be explained by an inter-division adder model, i.e. assuming that cells directly constrain the volume added between birth and division, other models also predict the adder phenotype. For example, even before the papers demonstrating the adder phenotype appeared, theoretical work by Amir showed that such an adder phenotype is also predicted by a model that combines an inter-initiation adder with a fixed time between replication and division, i.e. as in the Helmstetter-Cooper model (Amir, 2014; Ho and Amir, 2015; Amir, 2017).”*

Historically, Koppes and colleagues were the first to observe and analyze the adder phenotype in Voorn, Koppes, and Grover (1993). For example, they calculated correlations coefficients *r* between birth size and division size and obtained values close to *r*=1/2, and coined the model ‘incremental size model’, which is what we call the adder. The adder phenotype was proposed as early as 1973 (Sompayrac and Maaloe 1973), who predicted the “inter-initiation adder model” (Witz *et al.*’s terminology) using the balanced biosynthesis and threshold hypotheses. As for our own work, it suffices to say we “rediscovered” and documented the adder principle in early 2012. See also (Jun et al. 2018) for more information on the history of the adder principle.

*“in our work we thus stressed that, in order to meaningfully compare the evidence that the single-cell data provide for one or another model, one should compare the **full correlation structure** of the data with the predictions of the different models. We show in Witz et al. (2019) that, while the data are consistent with an inter-initiation adder model, the time between replication initiation and division in fact correlates with growth-rate (Fig. 3A), thereby falsifying a model that assumes division is controlled by a mechanism that acts on the time between initiation and division. Instead, our analysis shows that in the model which is most consistent with the full correlation structure of our data, division is controlled by an adder mechanism that runs from replication-initiation to division. In this model, each time replication is initiated an adder is started for each replicated origin, and each of these adders trigger a subsequent division event when a total critical amount of volume has been added. Importantly, we show that this model also reproduces the ‘adder phenotype’, i.e. that added volume between birth and division is approximately uncorrelated with the cell’s size at birth (Fig. 5A). Thus, the model that we propose assumes that an inter-initiation adder controls replication initiation, that division is controlled by an adder running from replication-initiation to division, and that growth-rate fluctuates independently from the replication and division control. The model proposed in the comment of Le Treut et al. is almost identical. First, the model of Le Treut et al. also assumes that replication initiation is controlled by an inter-initiation adder and in this sense replication control is thus ‘replication-centric’ in both models. Second, both models assume that growth-rate is an independent variable whose fluctuations are uncoupled from the control of the sizes at which replication and division occur. Third, both models also agree that division is controlled by an adder mechanism. The only difference is that, while our model assumes this adder is initiated at replication initiation, the model that Le Treut et al. propose assumes the adder is initiated at birth. For future reference we will refer to these models as the Replication Double Adder (RDA) and Independent Double Adder (IDA) models.*

### What does the evidence say?

*Although Le Treut et al claim that the IDA model has been ‘revealed’ in Si et al. (2019), a reader might be excused for not coming away with this impression from reading this paper. Si et al. (2019) show that cells exhibit ‘adder phenotypes’ for both the replication and division cycles, i.e. that the interinitiation added volume is approximately uncorrelated with the size at initiation, and that the added volume between birth and division is approximately uncorrelated with the size at birth. Then, by applying various perturbations, including massive time-dependent perturbations in the levels of key cell cycle proteins, Si et al. (2019) show that the correlations that define the replication adder and division adder phenotypes can be separately perturbed. **However, instead of putting forward the IDA model, in Si et al. (2019) the authors develop a complex variant of the Helmstetter-Cooper model which allows for an arbitrary correlation structure between all variables in the model, as well as correlation across generations, without discussing how these complex correlations could be implemented mechanistically.”***

Witz and colleagues appear to misunderstand Si et al. (2019)’s approach and analysis (e.g., “Also, that study focuses exclusively on a model which explicitly enforces various correlations between variables unlike our model which naturally produces such relations.”). The analysis in (Si et al. 2019) of the H-C model was to (1) falsify the stochastic version of the model and (2) point out the degeneracy; that is, even for the non-adder model, there is a significant fraction of parameter space that can fit the adder phenotype when ad-hoc fluctuations are introduced. Please refer to (Si et al. 2019) and (Facchetti et al. 2019) for further information.

*“In fact, we see no reason why any of the results in Si et al. (2019) would favor the IDA over the RDA model. Given that the replication and division cycles are ultimately implemented by mechanisms that involve separate molecular players, one would generally expect that it is possible to separately perturb the two adder phenotypes. In order to predict how either the IDA or RDA models would respond to the perturbations applied in Si et al. (2019) one would need much more concrete models of how these adder mechanisms are implemented biophysically. In any case, no attempt is made in Si et al. (2019) to evaluate to what extent their data favor one model over the “*

*“In contrast, **we did explicitly investigate whether the IDA model could also explain our data.** Simulations of the IDA model showed that, while this model reproduces most of the statistics observed in our data, it clearly fails in one critical respect (Appendix 2 Figure 2 of Witz et al. (2019)).”*

The ***I***-value analysis was never applied to the IDA model in (Witz, van Nimwegen, and Julou 2019), and only in their response they clarified that the ***I***-value analysis is not applicable to the IDA model. As pointed out earlier, they ruled out the IDA model based on one correlation from one dataset.

*“That is, **the IDA model predicts a strong negative correlation between the size at replication-initiation and the size added between initiation and division. However, no such correlation is observed in the data.** Even though we thus specifically identified a correlation for which the IDA and RDA models make different predictions, and showed that our data clearly favor the RDA over the IDA model, Le Treut completely ignore this observation, never comment on it, and make no attempt to investigate it on their own data. This is especially remarkable given that Le Treut et al. go as far as to claim, in direct contradiction to these results, that our own data in fact favor the IDA model over the RDA model. We next discuss why those claims are based on faulty analysis.”*

See the section “4. The RDA model is based on one dataset, but the full datasets show different results.”

### “The correlation analysis and independence measure I apply only when the events of each cell cycle are independent of other cell cycles

*In Witz et al. (2019) we developed a correlation analysis method to systematically compare a large class of cell cycle models on our largest, automatically curated, dataset, which was for cells growing in minimal media with glycerol. To rigorously compare how well different cell cycle models can explain a dataset, one would generally have to formulate concrete likelihood functions that calculate the probability of all the observed data given each model, i.e. all times and sizes at birth, replication initiation, and division, for all observed lineages of cells. Depending on the growth conditions and models one wants to consider, such likelihood functions will typically contain non-trivial dependencies between events across multiple cell cycles.*

*However, we took advantage of the fact that, in the growth conditions we studied, cell cycles do not overlap, and considered a class of models for which, given the state of the cell at the start of its cell cycle, all events in each cell cycle are independent of the events in other cell cycles. In particular, the class of models we considered are defined by the starting point of the cell cycle, which may be at replication-initiation or birth, and by a set of variables that are being independently constrained. For example, for the Helmstetter-Cooper model each cell cycle starts at replication-initiation and is defined by the growth-rate, the size at initiation, and the time between initiation and division. Similarly, in the RDA model the cell cycle also starts at initiation and is defined by growth-rate, the added volume between initiations, and the added volume between initiation and division. Given a model, the data of each cell cycle are defined by the state of the cell at the start of the cell cycle, and the values of the variables that the model assumes are independently constrained. As discussed in Witz et al. (2019), in this situation the fit of each model to the data can be quantified by the determinant I of the correlation matrix. We find that, of all models considered, the RDA model has the highest value of I and we subsequently showed, using simulations, that the RDA model also fits the data from experiments at higher growth rates.*

***However, as explained above, this correlation analysis crucially relies on the fact that cell cycles are not overlapping.** As soon as cells grow so fast that cell cycles overlap, events within one cell cycle depend on events that occurred one or more cell cycles in the past, and a more complex correlation analysis would be required. In addition, for the IDA model there are two adders running concurrently, so that there is no single event at which the cell cycle can be considered to start. **Consequently, the correlation analysis presented in our paper cannot be applied to the IDA model even at slow growth,** because events within one cell cycle depend on at least one event from a previous cell cycle. Indeed, exactly for this reason we did not attempt to calculate an I value for the IDA model, but instead highlighted one key correlation for which the RDA and IDA models make different predictions, as we mentioned above.*

***In hindsight we realize that we should have explained more clearly that the I values of our correlation analysis can only be meaningfully calculated when all cell cycles are non-overlapping and for models for which the events in each cell cycle are independent of the events in other cell cycles.”***

In this clarification, Witz *et al.* confirm that their ***I***-value analysis is only applicable to *E. coli* with non-overlapping cell cycles. This is unfortunate since the hallmark of *E. coli* cell cycle is its multifork replication. For example, the Helmstetter-Cooper model was directly motivated by the existence of overlapping cell cycles. Furthermore, the celebrated (nutrient) growth law (Schaechter, Maaløe, and Kjeldgaard 1958) is also applicable to the “fast” growth conditions wherein cell cycles typically overlap. By contrast, *E. coli* does not follow the adder principle in slow-growing cells as reported by Wallden et al. (2016) and explained by Si et al. (2019). This clarification, however, raises other questions, such as: Is the RDA model also limited to non-overlapping cell cycles? Moreover, since the ***I***-value analysis cannot be applied to the IDA and other models, the conclusion reached in (Witz, van Nimwegen, and Julou 2019), namely that initiation of chromosome replication controls both replication and division, is questionable.

***“Le Treut et al. have clearly not realized this limitation** and most of the results in their comment consist of comparing I values for the RDA and IDA models, including on datasets with overlapping cell cycles. However, since one cannot meaningfully calculate an _7_ value for the IDA model, nor for growth conditions with overlapping cell cycles, all these reported results are essentially meaningless.”*

It is the authors’ responsibility to state the limitations of their methods clearly. Apparently, none of the referees caught the limitations during the review process.

***“The correlation analysis does require 4 and not 3 variables***

*However, apart from these problems, the I values that Le Treut et al. report are also meaningless because of an additional error. **Instead of following our method and calculating the determinant of a 4 by 4 correlation matrix of 4 variables, Le Treut et al. claim that it suffices to calculate the determinant of only a 3 by 3 correlation matrix.** In particular, they argue this suffices because cell cycle models can be defined using 3 variables only. While it is true that 3 variables may be enough to define a cell cycle model, it is not enough to fully characterize the correlation structure of the cell cycles.*

*To illustrate this we will use a simplified example by ignoring the replication-cycle, and considering only the division cycle running from birth to division. The measured variables for a cell cycle will then consist of the length at birth Lb, the length at division Ldand the time T between birth and division. In our correlation analysis, each possible model describes the events of each cell cycle by the state of the cell at birth and two additional variables that control the timing and size at division, and then calculates the determinant I of the 3 by 3 correlation matrix of these variables. For example, the top row of Fig. 1 shows the correlation matrices and independence values I for the adder model and for a sizer model as calculated from our experimental data in slow growth conditions (M9-glycerol) used in Fig.6 of (Witz et al., 2019). That is, for the adder model each cell cycle is characterized by size at birth Lb, added length dL = Ld - Lb, and the growth-rate L Similarly, for a sizer model each cell cycle is characterized by size at birth Lb, size at division Ld, and the growth-rate L As expected, the I values indicate that the adder model is by far preferred over the sizer model.*

*However, one might argue that only 2 variables are needed to define the adder and sizer models, i.e. for each cell cycle we can draw the growth-rate λ and either dL or Ld from an independent distribution. However, as shown in the panels of the bottom row of Fig. 1, if we leave the initial state Lb out of the correlation matrix, and calculate I values for the 2 by 2 correlation matrices, there is no longer any meaningful difference between the I values of the two models. That is, the crucial evidence for the adder model is in the correlation between the sizes at birth and division and this is lost when the initial size Lb is left out of the correlation matrix. While it is true that one can think of the adder model as only specifying λ and dL of each cell cycle, once we leave out Lb the size at birth of one cell depends on the values of dL in all its ancestor cells, thereby coupling the data from different cell cycles together. Since the correlation analysis requires treating each cell cycle as independent, it is thus crucial to include the size at birth. “*

This is inaccurate, and calculations of the determinant of a 4×4 matrix of 4 variables had been available with (Le Treut et al. 2020). For full results, see the section “2. The *I*-value analysis in its current form cannot compare different models.

*“Unfortunately, Le Treut et al. have also not recognized this problem, and consistently calculate I values for 3 by 3 correlation matrices. These I values are thus meaningless for at least two separate reasons.”*

As mentioned earlier, it is the authors’ responsibility to state clearly any significant limitations of their methods for both the referees and the readers. Furthermore, as shown in the section “2. The ***I***-value analysis in its current form cannot compare different models”, both the 4-variable analysis and the 3-variable analysis have been available with (Le Treut et al. 2020), and their results are mostly consistent with each other.

### “Since Le Treut et al’s correlation analysis is invalid, discussions about the role of slow growth, the precise strain, or the origin tracking method are all moot

*Le Treut et al. spend a considerable portion of their comment comparing I values of the RDA and IDA models for datasets from different experiments, showing that the I values they calculate slightly prefer either the RDA or IDA model depending on whether one considers slow growth conditions, what precise E. coli lab strain is used, whether one supplements uracil in the growth media, and whether one uses FROS or dnaN-YPet to track replication initiation. Since, as we just explained, these I values result from a faulty analysis, the small differences that are observed are meaningless and thus not worthwhile discussing.*

*However, even if we were to assume that the I values were meaningful, the logic of Le Treut et al’s discussion of these results seems rather confused. **That is, Le Treut et al. suggest that the fact that RDA model is preferred over the IDA model on some of their own datasets** is an artefact resulting either because of slow growth, or because of a growth defect due to a rph-1 mutation that requires uracil supplementation to overcome, or due to using FROS. However, as we already mentioned above, **Le Treut et al. also claim that our own data prefer the IDA over the RDA model**, and those data were obtained using a strain with the same rph-1 mutation as strain MG1665, without supplementing uracil, and using FROS instead of dnaN-YPet. This thus directly contradicts that these details would cause the RDA model to be ‘artificially’ preferred.”*

This is inaccurate. In the 3-variable analysis, 15/19 datasets support the IDA model according to the ***I***-value analysis. In the 4-variable analysis, 12/19 datasets in both our study and Witz *et al.* support the IDA model, but with the uracil supplement in the media only 4/19 datasets support the RDA model (see next paragraph).

MG1655 is pyrimidine pseudo-auxotroph (Soupene et al. 2003) and, when supplied with uracil, the ***I***-values of the IDA model became larger than the ***I***-values of the RDA model. This indicates that the ***I***-value analysis is sensitive to the genetic background used in the experiments. As for the BW strains, to the best of our knowledge, it is unknown whether the BW strains also show pyrimidine auxotrophy regardless of *rph-1* mutation. However, the addition of uracil again reverted the ***I***-values in the 4-variable analysis of the BW strains data.

See Figure 2 in the section “2. The ***I***-value analysis in its current form cannot compare different models” for more details.

### “Independence values Ican only be meaningfully calculated for valid cell cycle decompositions

*Even though most of the results that Le Treut et al. present consist of comparison of /values, in the last part of their comment they go on to claim that these I values are in fact too insensitive to meaningfully distinguish between different cell cycle models. To support this claim, they start from their set of 18 possible variables that can be used in cell cycle decompositions and calculate I values for all 816 possible subsets of 3 variables, showing that these Ivalues have a smooth distribution, and noting also that the IDA and RDA models rank only 57th and 88th, respectively. **While we have some understanding for the previous mistakes made by Le Treut et al., i.e. mistakenly assuming that it was meaningful to calculate I values for the IDA model and in situations with overlapping cell cycles,** and mistakenly assuming that a 3 by 3 correlation matrix would suffice, we are frankly astonished that they do not seem to realize that the vast majority of these 816 subsets cannot specify cell cycle models at all. For example, size at birth L_b_, size at division L_a_, and added size d_L_ = L_d_ – L_b_ would be one of the 816 possible subsets. However, this subset does not specify anything about either replication initiation nor anything about the timing of the cell cycle events. It is of course completely meaningless to calculate I values for random subsets of variables like this. Moreover, we explicitly showed in our paper (Fig. 7A), that if one calculates I values only for meaningful decompositions (listed in Fig. 7S3), the top RDA decomposition clearly stands out, and all division-centric decompositions perform poorly. “*

Certainly, not all of the 816 combinations correspond to biologically realistic models of the cell cycle. However, the point of this analysis is to show that many such biologically unsound models can produce ***I***-values higher than biologically sound models. Since ***I***-values cannot distinguish sound from unsound models, the ***I***-value analysis lacks predictive power. See the section “2. The ***I-***value analysis in its current form cannot compare different models.” for more detail.

### “The RDA model does reproduce the observed ‘adder phenotype’

*Le Treut et al. also claim that the RDA model cannot reproduce the adder phenotype, i.e. that the added volume between birth and division is approximately independent of the volume at birth. To support this they analytically calculate the expected correlation coefficient between size at birth and division for an idealized version of the RDA model and show that this predicted correlation, while positive, is less than the correlation of 1/2 that would be predicted for a simple adder model. However, in our paper we explicitly showed through simulations of the RDA model that it does reproduce the ‘adder phenotype, i.e. see Fig. 5A of Witz et al. (2019).*

*So how is this possible? The reader will have to turn to the materials and methods of the comment of Le Treut et al. to discover that, although they used our simulation code to simulate the RDA model, they in fact modified the code so as to remove small stochastic fluctuations in the sizes of the two daughters at division. That is, we observed in the experimental data that a mother cell of size 2L does not produce two daughters of perfectly equal size L, but daughters of sizes L + δ and L-δ where δ/L has a standard-deviation of about 0.06, i.e. six percent fluctuations in the sizes of the two daughters. We incorporated this asymmetry into our simulations because it significantly improved the fit of the observed and predicted variation in cell size at birth, but it also improves the ‘adder phenotype’, i.e. the fit between the predicted and observed correlations in added volume dL and size at birth Lb of Fig. 5A. Indeed, in the Materials and Methods of their comment Le Treut et al. in fact admit that a mismatch between model and data is only observed when this asymmetry at division is removed.”*

The point of our theoretical analysis is not to state that the authors’ simulations do not agree with the experiment, but instead that the agreement between data and simulations critically relies on a fifth parameter, namely the variance in the division ratio σ_1/2_^2^. This additional variable was not used in the ***I***-value analysis and the RDA model. See the section “3. The RDA is analytically incompatible with the adder principle” for more detail.

*“We cannot understand why Le Treut et al. consider the fact that a simplified version of the RDA model (i.e. without asymmetry at division) predicts a correlation coefficient of less than 1/2 is considered an inherent flaw of the model. Whether the RDA model makes the exact same predictions as a perfect adder model is irrelevant. **The only thing that matters is whether the model fits the data.** Indeed, while the perfect adder predicts precisely zero dependence between the size at birth Lb and the added size dL, the RDA model predicts a weak negative correlation.**”***

Data fitting is not necessarily proof of a model’s validity. For example, the Helmstetter-Cooper model, with a combination of ad-hoc fluctuations and correlations, can fit the adder-type correlation data (Figure 2, (Si et al. 2019)).

*“In fact, a careful look at Fig. 5A of Witz et al. (2019) shows that the experimental data appear to exhibit a small negative correlation between Lb and dL as well. Moreover, such weak negative correlations can also be seen for the data shown in Fig. 3B and 4B of Si et al. (2019).*

*As noted earlier, several studies have shown that there is a clear positive correlation between size at birth L_b_ and size at initiation L_i_, (e.g. Fig 2A and 5B of Witz et al. (2019)), rejecting models that assume a critical initiation size, because such models predict no correlation between Lb and Li. It is thus noteworthy that a simple analytical calculation shows that, without asymmetry at division, the IDA would also predict no correlation between the size at birth L_b_ and size at initiation L_i_ However, when we simulated the IDA model including asymmetry at division, a weak positive correlation between L_b_ and Li appeared (Fig. 2 of Appendix 2 Witz et al. (2019)). Although this correlation is clearly weaker than what is observed in the data, we considered the difference too small to reject the IDA model on that basis, **and instead focused only on the clear mismatch in correlation between size at initiation and added size between initiation and division.”***

We discussed this in the Results section “4. The RDA model is based on one dataset, but the full datasets show different results.” Furthermore, one can calculate the correlation of the IDA model analytically and obtain:
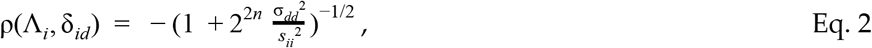

where σ_dd_^2^ is the variance of the birth-to-division added size Δ_d_, s_ii_^2^ is the variance of the unnormalized initiation-to-initiation adder Δ_ii_, and 2^n^ is the number of origins of replication at cell birth. Therefore, when the number of overlapping cell cycles *n* increases, the negative correlation in Eq. 2 approaches zero since the ratio σ_dd_^2^/s_ii_^2^ is of order 1 and does not change significantly under different nutrient limitations (Si et al. 2019; Taheri-Araghi et al. 2015).

*“Finally, we think it is instructive to compare the amount of concern that Le Treut et al. express regarding the small difference in the precise correlations predicted by the RDA and perfect adder models, with the concern for accuracy in such matters that these authors exhibit in Si et al. (2019). For example, whereas in Witz et al. (2019) we make sure to display experimentally observed correlations using full scatters, in Si et al. (2019) correlations are only shown using binned data, and these binned data are plotted using symbols that are so large that data from different experiments completely obscure each other, making it virtually impossible to see what precise correlations the data actually exhibit, e.g. see Figs. 1D and 7B of Si et al. (2019). Moreover, in Si et al. (2019) the authors routinely describe observed correlations as ‘sizers’ or ‘adders’ when the observed correlations clearly deviate substantially from what sizer and adder models would predict. For example, Fig 3B of Sietal. (2019) purports to show that, under the perturbations in question, the relationship between size at initiation and inter-initiation added size behaves as a ‘sizer’, while the division cycles still show adder behavior, i.e. that the added length between birth and division is independent of birth size. However, the negative correlation between inter-initiation added size and initiation size is clearly less steep then a sizer would predict, and there is also clearly a weak negative trend between size at birth and added size (as an RDA model would in fact predict). Very similar comments apply to Fig. 4B of Si et al. (2019) where the authors call a variety of clearly different slopes all ‘sizer’ whereas a clearly still negative but weaker slope is called an ‘adder’. The very weak negative correlation in Fig. 7B, which the authors call a ‘sizer, is also far from what a sizer would actually predict.”*

Si et al. (2019) provide separate scatter plots for all correlations in the supplementary information. Besides, the point of Figures 4B and 7B in (Le Treut et al. 2020) is that, once we understand the mechanisms underlying the adder, we are able to reprogram the size-control phenotype experimentally in a quantitatively predictive manner.

### “Conclusions”

*It has not been a pleasant exercise to write this response. We more than welcome post-publication review and feedback on our work, but we struggle to find anything constructive in the comment of Le Treut et al, and it is hard for us to understand what motivated these authors to attack our work so vehemently and unreasonably. This is especially surprising to us given that the IDA model that they champion in their comment is so close to the RDA model that we proposed in our paper. In fact, in Si et al. (2019) these authors repeatedly state that the main result of their experiments and analyses is that the division adder phenotype is a direct consequence of: (1) (threshold) accumulation of division initiators and precursors to a fixed threshold number per cell and (2) (balanced biosynthesis) their production is proportional to the growth of cell volume under steady-state condition. However, both of these points apply equally well to the initiation-to-division adder of the RDA model proposed in Witz et al. (2019). As such, we do not understand why Le Treut et al. even feel that the results presented in Witz et al. (2019) are at odds with theirs. That is, we do not see any result in Si et al. (2019) that suggests that the division adder must be initiated at birth rather than at replication-initiation.”*

The RDA and the IDA models represent fundamentally different biological mechanisms, regardless of the correlations.

*“Finally, we would like to stress that we do not believe that these relatively simple models give anywhere near a full picture of bacterial cell cycle control. On the contrary, it seems likely to us that the evolutionary process has ‘designed’ cell cycle controls that involve multiple complementary and redundant mechanisms including multiple check points. Moreover, different mechanisms may be more or less dominating in different species or within different conditions. For example, although both the IDA and RDA models do not specify any direct coupling between the replication and division adders, we of course know that couplings must exist. For example, we know that under particular stress conditions, such as when cells filament as a result of DNA damage, additional checkpoint mechanisms clearly come into play. In our simulations of the IDA model we observed a small fraction of cell cycles where the division adder was triggered before replication had even been initiated (let alone finished), and cells undoubtedly have control mechanisms to avoid this. Indeed, Micali et al. (2018b) recently introduced a family of models that combine concurrently running processes that separately control the replication and division cycles with an explicit check point mechanism and, as we pointed out in Witz et al. (2019), it is plausible that such models could in principle also fit our data, provided that their parameters are precisely tuned. In summary, we feel that models such as our RDA model should only be considered as useful starting points for further exploration.”*

The title of the original *eLife* paper is very clear and general, “Initiation of chromosome replication controls both division and replication cycles in *E. coli* through a double-adder mechanism.” It is therefore unfortunate that we are unable to reproduce the conclusions as elaborated in this report. At the same time, Witz *et al.*’s paper had the merit to motivate us to reassess various bacterial cell-cycle models, and we believe such an exercise and scientific exchanges can only benefit the field. We hope our report clarifies various subtle issues one may confront when obtaining conclusions from correlation analysis, and steer new experimental designs to test important hypotheses in this important field of quantitative microbial physiology.

